# Bioprinting transgenic plant cells for production of a recombinant biodefense agent

**DOI:** 10.1101/2021.02.01.429263

**Authors:** Anika Varma, Hawi B. Gemeda, Matthew J. McNulty, Karen A. McDonald, Somen Nandi, Jennifer M. Knipe

## Abstract

Transgenic rice cells (*Oryza sativa*) producing recombinant butyrylcholinesterase (BChE) as a prophylactic/therapeutic against organophosphate nerve agent poisoning, cocaine toxicity, and neurodegenerative diseases like Alzheimer’s were immobilized in a polyethylene glycol-based hydrogel. The cells were sustained for 14 days in the semi-solid matrix, undergoing a growth phase from days 0-6, a BChE production phase in sugar-free medium from days 6-12, and a growth/recovery phase from days 12-14. Throughout this period, the cells maintained similar viability to those in suspension cultures and displayed analogous sugar consumption trends. The rice cells in the bioprintable hydrogel also produced a significant amount of active BChE, comparable to the levels produced in liquid cultures. A considerable fraction of this BChE was secreted into the media, allowing for easier product separation. Overall, we demonstrate a simple, efficient, robust, modular, and potentially field-deployable bioreactor system for the manufacture of biologics.

## Introduction

Plant cell culture holds much promise in manufacturing foods [1], cosmetics [1], and biopharmaceuticals [2]. There are many benefits to plant cell culture over traditional cell culture, including cheaper and more readily available substrates and media, ability to perform complex post-translational modifications like glycosylation, and safety due to the inherent inability to propagate most mammalian viral contaminants [3–6]. On the other hand, plant cells in suspension cultures can be limited by slow growth rates, cell aggregation-limited mass transfer, and lower product accumulation [7]. Thus, we investigate a bioreactor design that involves immobilizing transgenic plant cells in a bioprintable hydrogel for the production of recombinant proteins. Bioprinting is an emerging process intensification strategy that provides a means for high cell loading, configurable geometries to facilitate mass transport of nutrients and products, and bioreactor versatility through the highly tunable and controllable hydrogel mechanical support [8–10]. This, ultimately, has the potential to increase reactor volumetric productivity and simplify recovery of secreted products [8].

Plant cell hydrogel immobilization is not necessarily a new idea; the concept emerged in the late 1970 **[11]**. Literature spans casting plant-encapsulated hydrogels, 3D bioprinting plant cell laden hydrogels, and crafting hydrogel-suspension culture hybrid systems for a variety of plant types including *Valerianella locusta* (lettuce) **[12]**, *Nicotiana tabacum* (tobacco) **[13]**, *Ocimum basilicum* (basil) **[14]**, *Taxus baccata* (English yew) **[15]**, *Taxus cuspidata* (Japanese yew) **[16]**, *Chlamydomonas reinhardtii* and *Chlorella sorokiniana* (microalgae) **[17,18]**, *Capsicum frutescens* (Tobasco pepper) **[19]**, *Solanum aviculare* (New Zealand nightshade) **[20]**, *Solanum malacoxylon* (Waxyleaf nightshade) **[21]**, and *Zinnia elegans* (common zinnia) **[22]**. Plant cell hydrogel immobilization has been demonstrated as an effective means to culture cells and improve cell culture productivity for natural secondary metabolite products like scopolin **[13]**, paclitaxel **[15,16]**, baccatin III **[15]**, capsaicinoids **[19]**, and choleocalciferol **[21]**. However, there are no reports in literature to date that demonstrate plant cell hydrogel immobilization as a strategy to enable mobile and field-deployable bioreactor systems to produce protein therapeutics, or other high-value proteins, which is the goal of this study.

As a model system, we employ transgenic rice cells from an optimized stable cell line to produce active butyrylcholinesterase (BChE), a native human enzyme mainly found in the blood plasma that functions as a bioscavenger against various organophosphorus nerve agents and has potential therapeutic application in Alzheimer’s disease [23–26]. Although current commercial preparations of BChE are purified from human (and animal) blood plasma, the high cost and unreliability of a plasma-derived supply chain advocates a recombinant source as a supplement or replacement [3]. Researchers have pursued recombinant BChE production in a range of expression systems including transgenic animals, such as goats and mice [27], whole plants, like *Nicotiana benthamiana* [28,29], and cell culture, including insect cells [30], CHO cells [31,32], HEK cells [33], and plant cells [34] with various degrees of success. The use of transgenic goats as bioreactors has resulted in production, purification, and formulation of a PEGylated form of recombinant human BChE from goat milk, and the drug that was developed made it into first-in-human clinical trials [35]. However, transgenic animal technology is extremely expensive and time-consuming to establish and maintain. Plant and CHO cell culture, which are significantly less expensive, have also demonstrated great success in producing active BChE, largely overcoming the difficulties in producing this complex tetramerized glycoprotein in a cost-effective manner [3,36–38]. Given the benefits of BChE production in plant cell culture, we aim to further develop the platform in a specialized bioreactor format, using hydrogel immobilization, to potentially serve as a rapid response, easily scaled out, and field-deployable tool to produce this biodefense agent for distributed rural and military applications.

Here we present our foundational work on hydrogel immobilization and bioprinting with transgenic *Oryza sativa* rice cells expressing BChE. We investigate hydrogel and bioprinting biocompatibility, impact on nutrient uptake, and recombinant BChE protein production. Because this study is primarily focused on bulk hydrogel material, there is substantial room for improvement of mass transfer and product recovery by tuning the cell-loaded hydrogel formulation, geometry, and reactor design. We explore the curing of bulk hydrogel disks as well as extrusion printing, which enables geometries and configurations that may lead to better mass transfer of gases, electrolytes, nutrients, and metabolites to the cells and protein product out to the media. This study demonstrates the feasibility of hydrogel-immobilized transgenic plant cells as a novel and promising bioreactor format to make biopharmaceutical proteins. This is a “proof-of-concept” study for the design of 3D printed plant cell bioreactors for continuous production of high-value recombinant proteins. Future work will build on this baseline to explore the cell immobilization and bioprinting design space, such as cell aggregate size distribution, cell loading, hydrogel geometry and composition, and optimize performance for sustained operation and specific applications.

## Results

In this study, we wanted to compare the viability and productivity of transgenic rice cells in suspension with and without UV exposure to those immobilized in PEGTA-LAP hydrogel disks. The immobilized cell condition is displayed in Figure 1, where a single disk was cut into 3 pieces.

**Figure 1.**
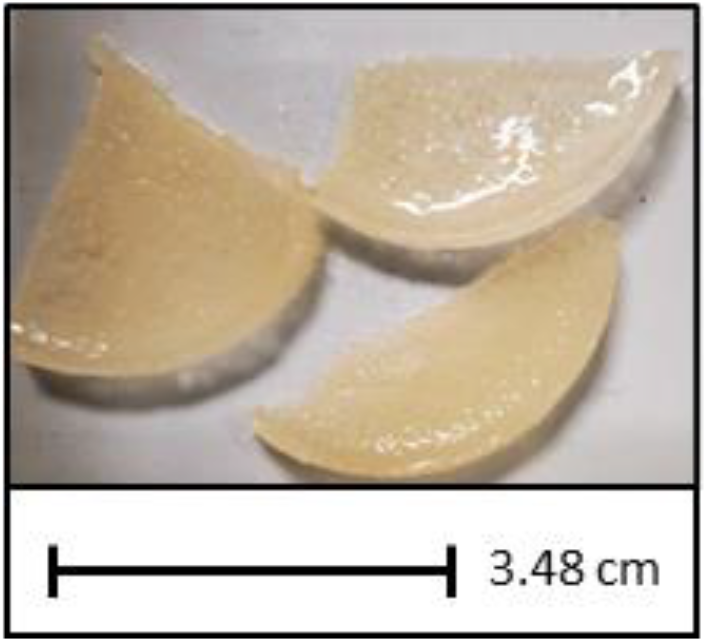
Cell-laden hydrogel. Representative image of UV-cured 12% (w/v) PEGTA, 0.1-0.2% (w/v) LAP in NB+S medium hydrogel disks encapsulating transgenic rrBChE-expressing rice cells at 50% (w/v), at a 2 mL working volume scale.

The first key outcome from the experiments performed here was cell viability. It was crucial that the transgenic rice cells remained viable in the hydrogel before we even considered rrBChE production and longevity of cells through multiple growth-expression cycles. The cell viability results are qualitatively displayed in Figure 2A, which shows TTC assay images. The darker the red color, the greater the metabolic activity, and thus inferred viability, of the cells. For all of the conditions, the viability was constantly high from days 0-6. By day 12, the cells appeared stressed (seen by the lighter coloration) due to sugar deprivation. Day 14 results indicated recovery of cell health. Additionally, the images suggest that the cells in the hydrogel may have been more metabolically active than in the control conditions on days 12 and 14. Figure 2B shows the quantitative TTC assay absorbance results. As correlated with Figure 2A, absorbance values and cell viability were high through day 6 and then dropped after induction of recombinant protein expression in sugar-free medium. The cells recovered when put back in growth medium, with the cells in the hydrogel experiencing less overall decline in viability. The trends in live/dead ratio as measured by fluorescence assay, exhibited in Figure 2C, further suggest that the fluctuations in cell viability are relatively small for all cell conditions through the duration of the experiment, although we acknowledge that propidium iodide stains cell walls in addition to the nuclei of dead cells, which may result in the ratio appearing artificially constant [39], as also seen in a calibration curve with known live/dead cell ratios (see supplemental figure S1). The live/dead ratio in the hydrogel sample at Day 12 is unexpectedly high compared with the TTC trend and may be due to greater background signal from the hydrogel in those particular samples. Confocal images of the suspension cells and hydrogel immobilized cells are shown in Figures 2D and 2E, respectively, where green is a nucleic acid stain and red is a cell wall and dead cell nuclei stain.

**Figure 2.**
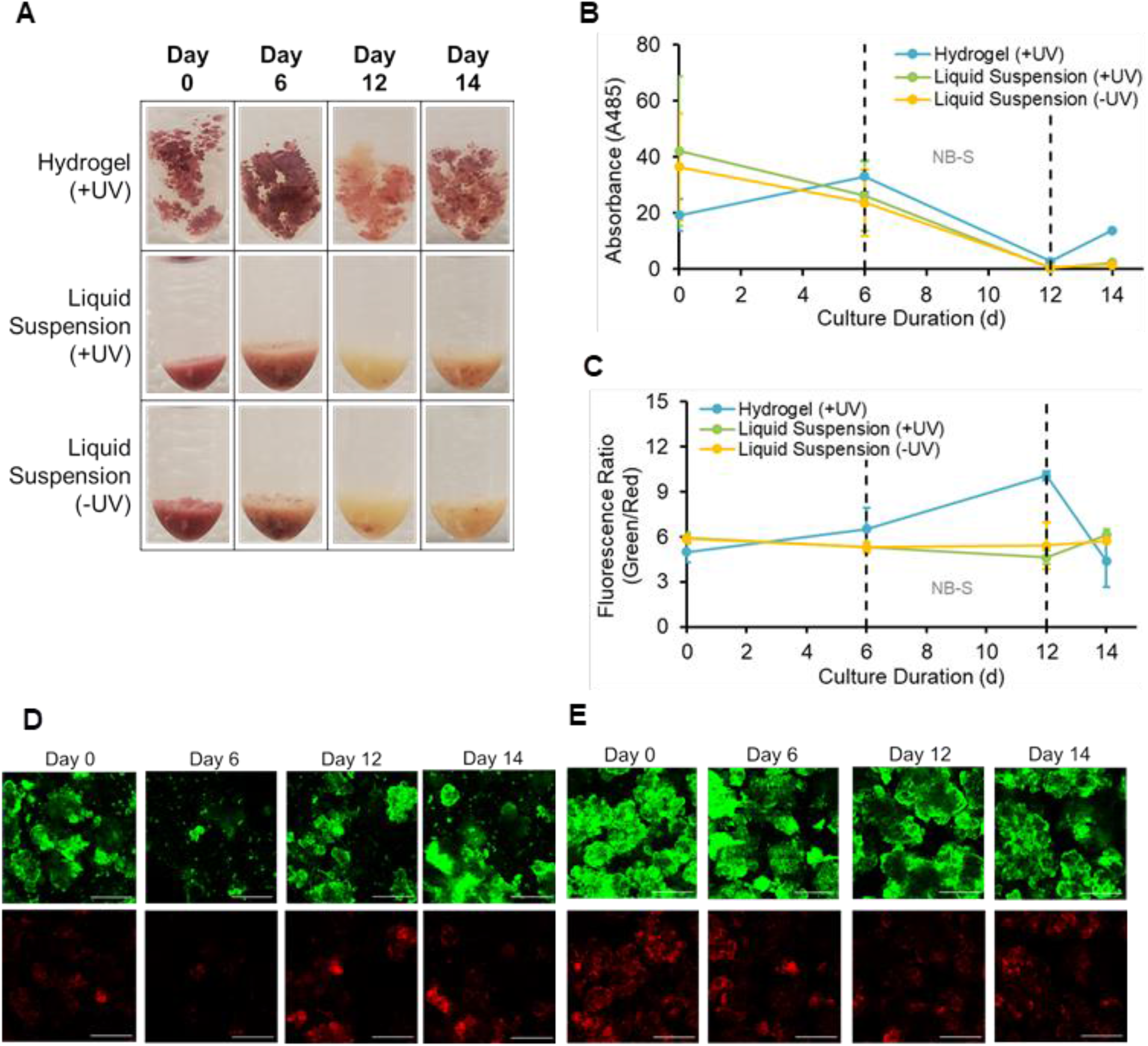
Viability of transgenic rice cells in hydrogel and in suspension. Transgenic rice cell viability over the course of 14-day culture (with media exchanges from sugar-rich to sugar-free medium on day 6 and from sugar-free to sugar-rich medium on day 12) as shown by (A) pictures of TTC-stained cells, (B) TTC assay absorbance readings, (C) live/dead cell ratio as calculated from fluorescent signal using plate reader, (D) dual-stained cells in suspension -UV under confocal microscope (200 μm scalebar), (E) dual-stained cells in hydrogel +UV under confocal microscope (200 μm scalebar). Error bars represent ± 1 standard deviation from biological duplicates and technical triplicates. Day 14 measurement of cells in hydrogel (+UV) for the TTC absorbance in (A) and (B) was based on a single biological replicate.

Figure 3 displays the sugar levels in each of the rice cell suspensions/hydrogels. Sucrose levels started close to the initial sucrose level in the media of 21.0 g/L and decreased steadily while glucose levels increased from 1 g/L to 4-8 g/L from days 0-6. After media exchange for induction of expression on day 6, sucrose and glucose levels dropped to 0, as expected, and remained there for days 6-12. After media exchange for recovery on day 12, sucrose and glucose levels were again at the maximum levels as prepared in the media. The trend for days 12-14 followed a similar pattern as for days 0-6, albeit with slightly steeper sucrose consumption and glucose production from the previously starved cells. Figures 3A, 3B, and 3C also indicate that sugar consumption was approximately equivalent among the cell conditions, implying that the cells in the hydrogel were just as metabolically active as those in suspension.

**Figure 3.**
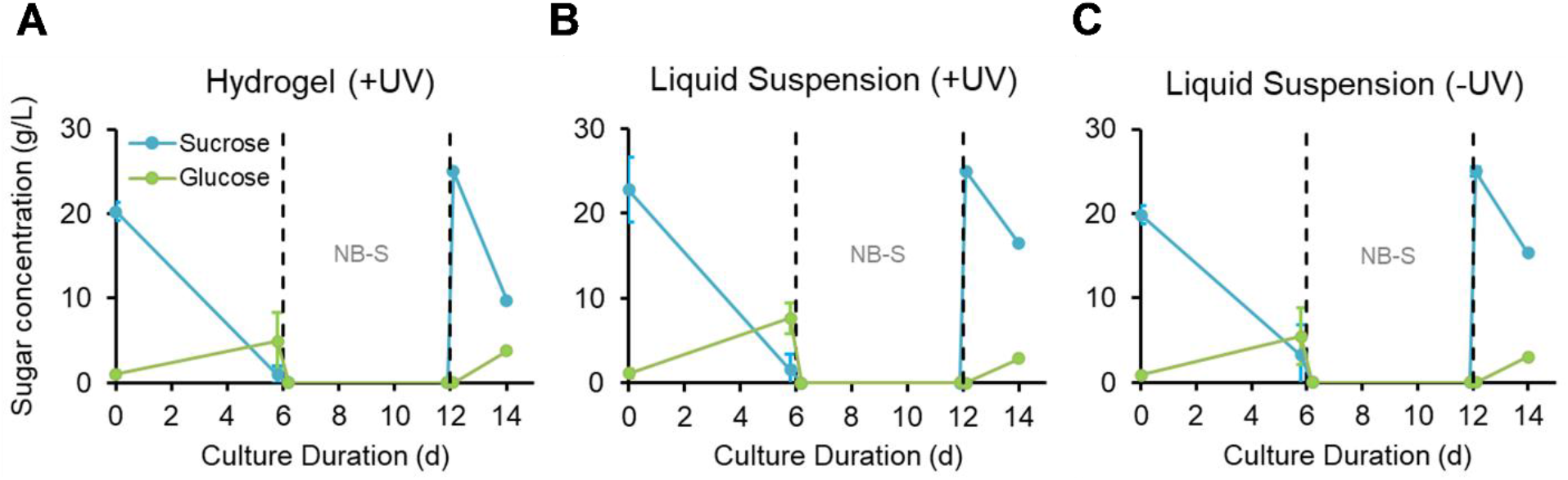
Sugar consumption of transgenic rice cells in hydrogel and in suspension. Sugar consumption trends over the course of the 14-day rice cell culture with cells (A) immobilized in hydrogel, (B) in liquid suspension with UV treatment, and (C) in liquid suspension without UV treatment. The dotted lines represent media exchanges; day 6 was the exchange from sugar-rich medium to sugar-free medium to induce rrBChE expression; day 12 was the exchange from sugar-free to sugar-rich medium to re-renter growth phase. Error bars represent ± 1 standard deviation from biological duplicates and technical triplicates. Day 14 measurement of cells in hydrogel (+UV) was based on a single biological replicate.

To investigate whether production of rrBChE was affected by the immobilization process we measured the levels of active enzyme on all sampling days. The levels were close to 0 on days 0 and 6, peaked on day 12, and were negligible in the media but still high in the cell extract on day 14. Figure 4 displays the rrBChE production levels at the end of the expression phase, on day 12. Figures 4A, 4C, and 4E are from one biological sample, and Figures 4B, 4D, and 4F are from a biological replicate, with error bars from technical triplicates. In all three conditions (hydrogel and liquid suspension controls), there was a significant quantity of rrBChE in the cell extract and also in the media, which was markedly different from previous work that used the same transgenic rice cell line [38]. This observation can be, in part, attributed to the lower biomass density (~14% w/v) and larger sieve mesh size used to reduce initial aggregate size distribution (380 μm) in the Macharoen et al., 2020 study [38]. In Figures 4A and 4B, the μg/g FW of rrBChE was defined as the total mass of rrBChE in the culture or hydrogel on day 12 normalized by the total initial g FW of cells at day 0. The quantity of media-secreted rrBChE was greater than the cell-associated across both biological replicates and conditions, barring the results of one of the controls (liquid suspension -UV) in Figure 4B. Figures 4C and 4D are similar to 4A and 4B but show the amount of rrBChE normalized by culture volume instead of by cell mass. In Figures 4C and 4D, the μg/mL of rrBChE was defined as the total quantity of rrBChE in a media or cell extract sample normalized by the respective volumetric compartment. For the media sample, this volume was ~10mL for both the liquid cultures and the hydrogel condition (which contained ~9mL liquid and 1mL solid). For the cell extract, this volume was the 1mL extraction buffer volume plus 1 g FW cells, for a total of ~2mL. With this measure, the rrBChE in the cell extract was slightly more concentrated than that in the media. Figures 4E and 4F depict the “starting” purity of active rrBChE in the media or in the cell extracts (no purification performed), computed as the percentage of TSP. The starting purities for the media and cell extract were consistently between 1.9-3.1%, no matter if the cells were immobilized or in suspension with/without UV curing.

**Figure 4.**
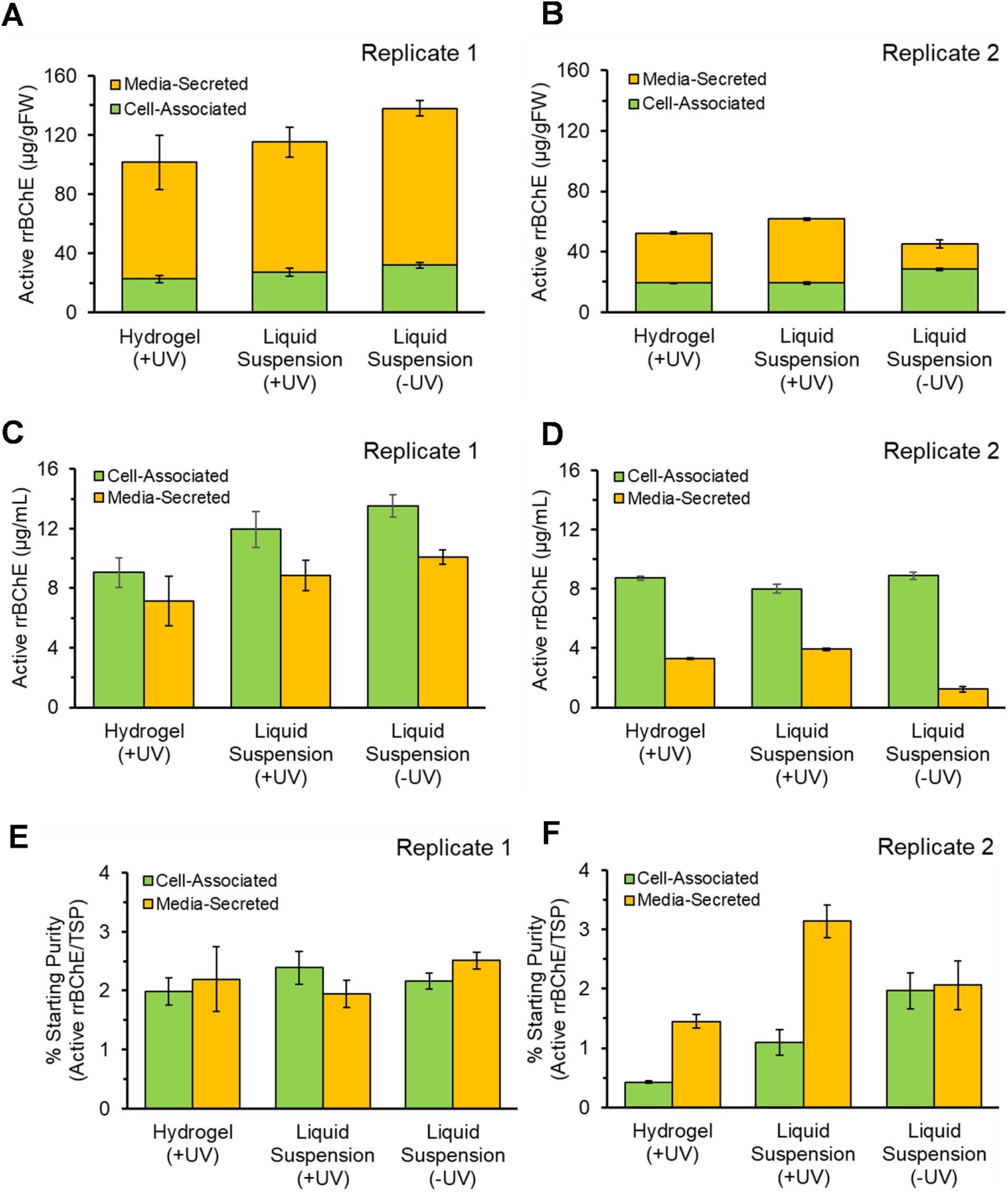
Active rrBChE levels for transgenic rice cells in hydrogel and in suspension. Profiles of cell-associated and media-secreted rrBChE concentrations on day 12 at the end of expression phase in sugar-free medium, separated by biological replicate: (A, B) rrBChE accumulation normalized by initial rice fresh weight; (C, D) rrBChE accumulation normalized by compartment volume, either cell extract or media; (E, F) starting purity of rrBChE normalized by TSP concentration. Error bars represent ± 1 standard deviation from technical triplicates. Day 14 measurement of cells in hydrogel (+UV) was based on a single biological replicate.

We qualitatively assessed the host cell protein profile using SDS-PAGE. Western Blot was carried out to confirm the presence and expected monomer size of rrBChE. This also visualized relative BChE production levels, ascertained quantitatively from the Ellman assay, between cells in the hydrogel and cells in liquid suspension with/without UV exposure. The gel and blot are shown in Figure 5. There was no discernible difference in the SDS-PAGE protein profiles between cell conditions, and Western Blot analysis confirmed the presence of rrBChE at the expected molecular weight for the monomer subunit without a human signal peptide (~65 kD) (full blot not shown). The most prominent band on the SDS-PAGE gel was α-amylase (~45 kD), which is co-regulated with rrBChE by the RAmy3D promoter [40,41]. rrBChE was not clearly observed on the gel, except in the 500 ng control lane where there was a lack of host cell proteins, but it manifested distinctly on the Western Blot. As expected, there was no rrBChE in the day 0 or day 6 samples, but a clear band was present in the day 12 cell extract and media samples, after the rice cells had been induced by sugar starvation. The substantial quantity of cell-associated rrBChE, despite inclusion of a secretion signal peptide, was not surprising given the large tetrameric rrBChE size (~260 kD without glycosylation). This could be due to detecting rrBChE in transit through the secretory pathway, or limitations in mass transfer out of the cell aggregates as well as diffusion limitations for transport of rrBChE out of the hydrogel and into the media.

**Figure 5.**
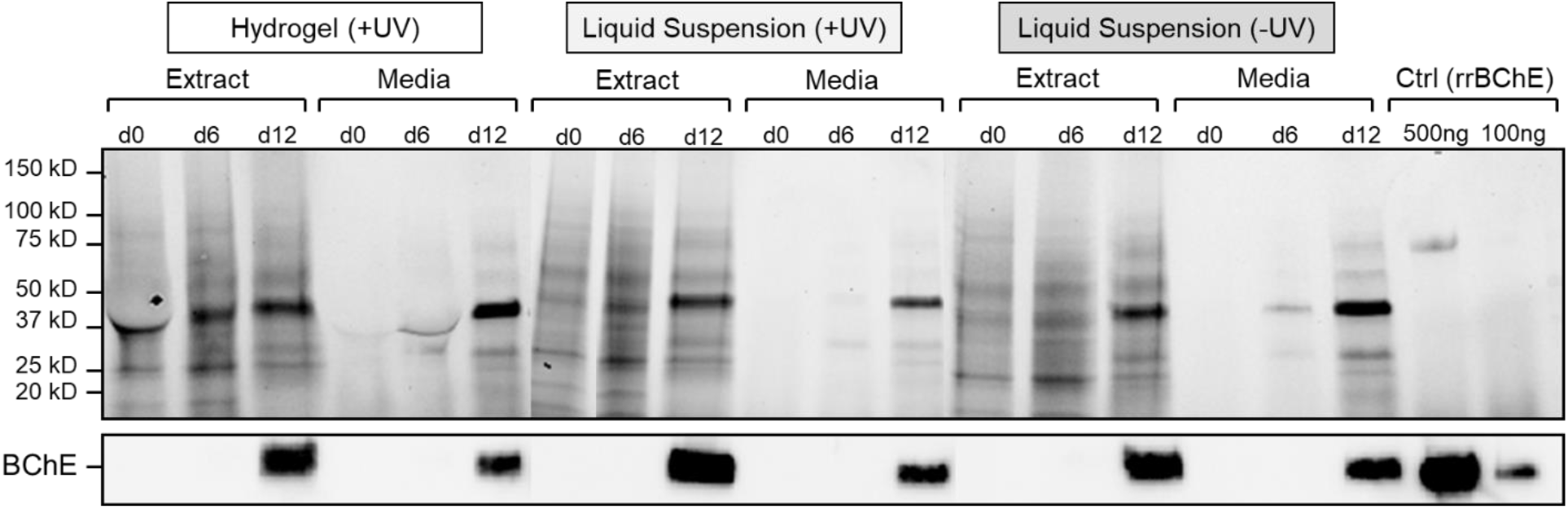
Protein profile and rrBChE band for media and cell extract samples from immobilized cells and suspension cultures. SDS-PAGE (top) and Western Blot (bottom) results displaying total protein profiles for cells in hydrogel, cells in liquid suspension with UV treatment, and cells in liquid suspension without UV treatment. The gel and blot depict both crude extract and culture media for each cell condition on days 0, 6, 12. Extracts were loaded to target 20-30 μg TSP, and media was loaded at 0-10 μg TSP. This corresponded to 475-625 ng rrBChE loaded in the day 12 extract lanes and 185-265 ng rrBChE loaded in the day 12 media lanes, as calculated from the active BChE Ellman assay measurements.

## Discussion

Transgenic rice cells can be successfully maintained in a PEGTA+LAP hydrogel with the same, if not higher, viability as in suspension culture, as supported by both TTC assay and confocal microscopy results. As mentioned in previous reports [8], the high 405 nm wavelength energy source proved biocompatible as we saw no difference in viability between cell suspensions exposed and unexposed to the UV lamp, shown in Figure 2. The cell viability results communicate that while the cells in the hydrogel are less viable on day 0, they recover and have slightly higher viability on days 6, 12, and 14 than either of the liquid suspension cultures (+UV or –UV) (see Figure 2). The lower viability of cells in hydrogel on day 0 could be due to the transient detrimental effects of the UV curing process. UV light induces the formation of free radicals in LAP, which cause oxidative stress on the cells and temporarily stall their growth and metabolism [42]. This effect is short-lived, however, and the cells recover by day 6. This would explain why only the hydrogel (+UV) condition experienced low viability on day 0. A hypothesis for the higher viability on days 6, 12, and 14 is that the immobilized cells sustained a slower growth rate than those in suspension. This was observed qualitatively by the relatively constant cell biomass in the hydrogel while the biomass for the liquid cultures noticeably increased through the 14-day experiment duration, though quantitative measurements of g FW were not taken. If the cells in the hydrogel were simply maintaining themselves instead of actively growing and dividing, the fewer number of cells would be able to support themselves with the same amount of sucrose (see Figure 3), and each cell would be afforded a higher metabolism. In the future, more sampling points between days 0-6 could confirm if this was the case.

In addition to maintaining viability, the cells in this study expressed and secreted commensurate amounts of total active rrBChE while immobilized in the hydrogel as while suspended in liquid culture, despite the high variability between replicate samples (see Figure 4). The variability can likely be attributed to variations in the initial biomass and the health of the cells during the experiments. When the experiments were originally set up, the initial cell biomass for each flask was targeted to be 1 g FW but measurements of actual initial cell biomass ranged from 0.76-1.15 g FW. This irregularity in biomass may explain some of the variability in culture performance, as more cells mean faster cell growth and sugar consumption, and the variability in rrBChE (μg/mL) concentration, as more cells mean higher rrBChE levels in the same liquid volume. Moreover, although the same cell line was used for two separate experiments, most of the cells were several weeks older during the second experiment and were not freshly subcultured from plates as the cells from the first experiment had been. The first experiment used 15-day old cells (from time of original plate-to-shake flask subculture), which were on their 2^nd^ passage in NB+S media, while the second experiment used primarily 33-day old cells on their 4^th^ passage in NB+S media with a smaller fraction of 13-day old cells on their 2^nd^ passage in NB+S media. Because the cells were, on average, much older for the second experiment, they may not have been as healthy and robust to begin with, which is supported by the variability in TTC viability data between the experiments (see supplemental figure S2). As the viability and likely the viable cell density of the cells was lower in the second experiment, the amount of rrBChE produced was also less. Furthermore, we postulate that the cell aggregate sizes were larger in the second experiment because cells were re-sieved earlier before the start of the experiment and, therefore had more time to recover and grow in size during the days prior to and during the start of the second experiment compared to the first experiment. These larger aggregate sizes could have meant that a larger percentage of rrBChE protein was retained within the aggregates rather than getting secreted, and the cells in large aggregates may have been less metabolically active per g FW, which is again reinforced by the difference in TTC results between experiments. Rather than focusing on the biological variability of the two experiments, we emphasize that the trends between cell conditions (hydrogel vs. suspension cultures with/without UV curing) are consistent and reproducible in both experiments. Namely, the rrBChE produced, whether in terms of μg/g FW or μg/mL, is comparable among all three cell conditions, and the starting purities are approximately equivalent. Therefore, we see that cell immobilization does not significantly affect recombinant protein production and secretion of our target molecule.

The positive viability and rrBChE productivity results drawn in this study incentivize continued scale-up and experimentation with cells immobilized in different geometries. This study was limited in scope to bulk hydrogel material, so there is still a lot of room for optimization of mass transfer and product recovery by manufacturing the hydrogel in various geometries. For instance, extrusion-based bioprinting, shown schematically in Figure 6A, may enable novel geometries and reactor designs. We have demonstrated a formulation of cells that can be extruded into a free-standing design, shown in Figure 6B, and subsequently cured. Bioprinted lattice structures with thin filament diameters may enhance mass transfer of nutrients into the hydrogel and rrBChE out of the hydrogel.

**Figure 6.**
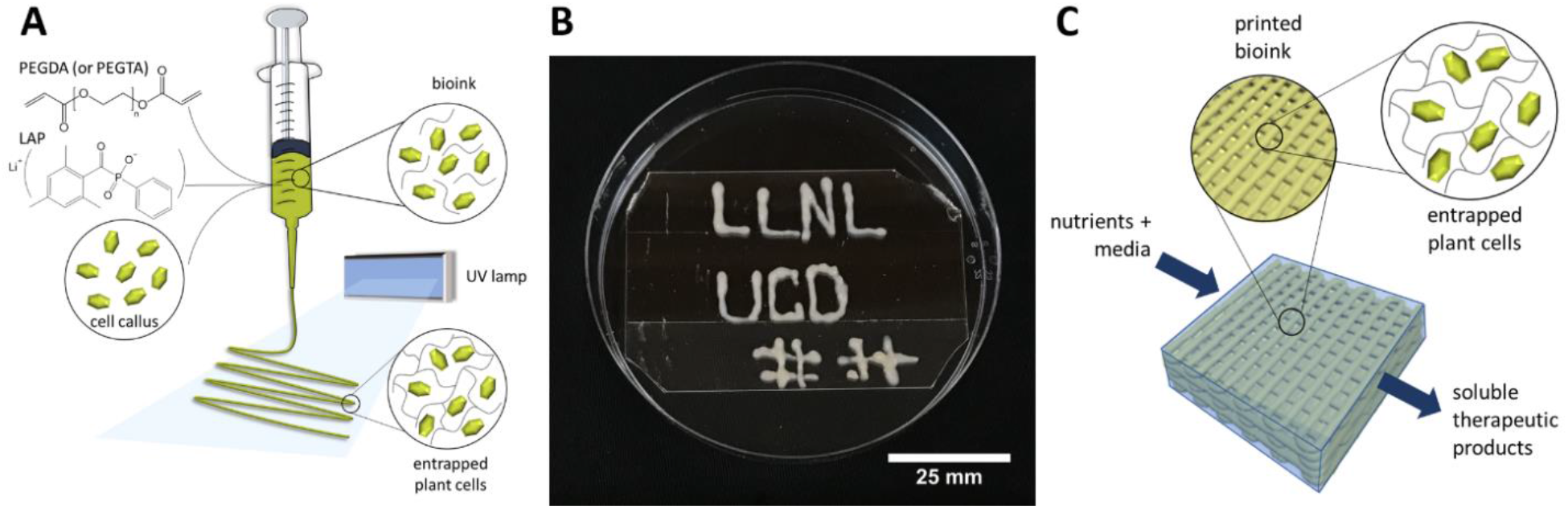
3D Bioprinting Plant Cells. (A) Transgenic rice cells can be immobilized in more complex geometries such as bioprinted lattices. (B) Initial extrusion results indicate a formulation with appropriate viscosity and high cell density. (C) Diagram of a flow-through reactor with entrapped, bioprinted plant cells.

Furthermore, in this study, a significant fraction of rrBChE was secreted into the media and was easily detected on a Western Blot, motivating the development of a flow-through bioreactor system, as illustrated in Figure 6C. In this system, immobilized cells could be maintained through multiple growth-expression cycles, and media exchanges would be very simple with enhanced sterility since no centrifugation or settling of cells would be necessary. Media from the expression phase could be recirculated and ultimately pumped out of the system to be ready for immediate purification and isolation of rrBChE product, while fresh, sterile, oxygenated growth media could be pumped into and recirculated through the reactor during the growth/rejuvenation phase. Additional considerations of the impact of cell growth on nutrient transfer will be important for more complex geometries and multiple use cycles. For example, in the flow-through reactor concept illustrated here, two factors to study are the transient profile of nutrient flux to accommodate a growing cell population and the manner in which that population may impede mass transfer through the bioprinted lattices if the aggregates affect the macroporosity or diffusion coefficient of the hydrogel matrix.

The most inspiring aspect of this research is that the results have a much broader application beyond expression of rrBChE in rice cells. The ability to 3D-print a plant cell bioink for use in a continuous flow-through bioreactor configuration offers a production platform for numerous therapeutics with the potential to overcome the limitations associated with a diversity of current plant cell culture processes. Typically, plant cells are cultured in suspension in stirred, air-sparged bioreactors [4,43–46] or in a “packed bed” of plant cells using a membrane bioreactor [47], and in these conditions have been shown to maintain viability and productivity for several months. However, neither of these bioreactor systems allow precise control over the 3D position of the plant cell aggregates, cell aggregate size distribution, or nutrient and product concentration profiles, which can affect productivity, ease of product recovery, and product quality. New and radical bioreactor designs and configurations, as presented above, fabricated by 3D bioprinting plant cells may enhance mass transfer; control cell aggregation; immobilize the cells for reuse over multiple production cycles; and enable novel use cases for bioproduction, biotransformations, and bioremediation.

This work demonstrates the feasibility of immobilization of plant cells into hydrogel disks to serve as an efficient bioreactor system with a small footprint from which valuable protein prophylactics and therapeutics can be derived.

In this proof-of-concept study, we demonstrated the successful encapsulation of rrBChE-producing rice cells within hydrogel disks by verifying the viability and productivity of the immobilized cells. Thoughts on future work include performing longer-term and larger-scale continuous experiments, on the order of months, to study the viability, growth, and rrBChE production of rice cells through multiple growth-expression cycles. It will be especially important to investigate how to maintain cell viability from cycle to cycle to achieve prolonged use of the bioreactor system and sustained protein production. In the experiments presented here, a relatively high seeding density of cells was employed, but tests with different proportions of cells to bioink may prove a different cell loading to be ideal. It is also unknown whether cells immobilized in hydrogel experience a growth phase or maintain a stationary phase. We hypothesize that the fraction of rrBChE secreted to the media is dependent on cell aggregate size; in the future we would like to explore whether immobilization within the hydrogel can control the cell aggregate size to maximize protein secretion.

Configuring cells within a hydrogel or bioprinted matrix may offer several advantages toward process intensification, including high cell density, control of cell aggregates, secretion and recovery of concentrated protein product, and continuous production. Extrapolated benefits are numerous, including the simplicity of the system without need for constant attention and extensive utilities like agitation, aeration, heating, and cooling; compactness and field deployability, on-demand production of therapeutic compounds as needed, and decentralized production through modular design with multiple small units as opposed to scale-up of capital-intensive equipment.

With this work as a foundation, the focus going forward can be given to scaffolding and/or extrusion printing cells in various geometries to optimize manufacture, isolation of desired proteins from the hydrogel, and integrated bioreactor system designs, paving the way to a rapid and adaptable system for biotherapeutic production.

## Materials and Methods

### Rice Cell Line and Suspension Culture Preparation

A previously reported transgenic *O. sativa* rice cell line expressing recombinant rice butyrylcholinesterase (rrBChE) was used in this study [3]. Expression of rrBChE is controlled through the metabolically regulated rice alpha amylase 3D (RAmy3D) promoter, which induces protein expression under sugar starvation [38]. This allows for cyclical or semi-continuous operation of cell culture, in which cells alternate between a growth phase in sugar-rich medium and an expression phase in sugar-free medium. The RAmy3D system also contains a signal peptide that tags proteins for secretion [3]. This permits easy separation of the desired protein to pass on to downstream purification and enables retention and reuse of the plant cells.

Rice cells used in hydrogel immobilization experiments were cultured and maintained according to previously described procedures on semi-solid “NB+S” selection medium containing N6 macronutrients, B5 micronutrients and vitamins, 30 g/L sucrose, 1.8 g/L Gelzan™, 300 mg/L casein hydrolysate, 250 mg/L L-glutamine, 250 mg/L L-proline, 2 mg/L 2,4-dichlorophenoxyacetic acid (2,4-D), 0.02 mg/L kinetin, and 50 mg/L geneticin as the selection antibiotic [3,38].

Before the start of experiments, the cells were subcultured into liquid sterile NB+S medium (without Gelzan™ and geneticin) by pressing and sieving the calli through a sterile, stainless-steel, 280 μm mesh sieve to achieve small and consistent cell aggregates [38]. Suspension cultures were grown in 500 mL to 1 L shake flasks with a 20% working volume, and they were incubated at 28 °C, 140 rpm in a 19-mm circular orbit, in the dark. Media exchanges with fresh liquid NB+S medium were completed weekly to keep the cells nourished. When desired, transgenic cells were induced to produce BChE via sugar starvation, in which the sugar-rich NB+S growth medium was exchanged for sugar-free NB-S expression medium, containing the same components as NB+S medium but with no sucrose. The cells were kept in this sugar-free state for 6 days and then passaged back to NB+S growth medium for recovery. Cultures were sustained like this for up to 3 months before fresh semi-solid-to-liquid subculturing was repeated.

### Bioink Formulation

The bioink used to entrap the rice cells consisted of 12% (w/v) of 4-arm polyethylene glycol tetra-acrylate MW 20,000 (PEGTA) (4arm-PEG-ACRL, MW 20kDa, Laysan Bio, Arab, AL) as the bioink base, and 0.1% (w/v) of lithium phenyl-2,4,6-trimethylbenzoylphosphinate (LAP) (TPO-Li, CPS Polymers, Boulder, CO) as the photo-initiator [8]. To prepare the bioink, a 10x LAP stock solution was made in NB+S medium. The solution was vortexed vigorously to dissolve the LAP. The PEGTA solution was made separately in NB+S medium and also vortexed. Immediately prior to the start of experiments, LAP stock solution was added to the PEGTA solution, and the mixture was vortexed again. For extrusion printing, the bioink formulation was prepared by adding nanocellulose crystal powder (The University of Maine, Process Development Center, Orono, Maine) to a loading of 16 wt% with a solution of 12% (w/v) PEGTA, 50 wt% cells, and 0.1% (w/v) LAP. The bioink was extruded manually using a 1 mL sterile luer-lock syringe fitted with a ~840 um inner diameter tapered tip (Nordson EFD, Westlake, OH) and cured following the procedure below.

### Experiment Setup and UV Curing

Experiments were performed in a biosafety cabinet. Rice cells in suspension culture were re-sieved through a 280 μm mesh filter and passaged to fresh growth medium in a shake flask a few days before the start of the experiment to reduce aggregation and ensure healthy, exponential phase, growing cells. To begin the experiment on day 0, 10-15 mL cell culture samples were taken from the flask, and cells were allowed to settle by gravity in the 15 mL Falcon tube (no centrifugation necessary) before media was removed. Cells measured as grams fresh weight (g FW), were then added to the prepared bioink solution, or to fresh NB+S growth medium in the case of the liquid suspension culture controls, to achieve a 50% (w/v) cell loading density.

Cell-laden PEGTA-LAP bioink samples were UV cured within a 6-well plate (3.48 cm diameter, 0.21 cm height, 2 mL total volume) through exposure to high intensity UV light for 10 seconds, using the Loctite EQ CL30 LED 405nm Flood System with UV LED Flood Controller (Loctite, Henkel Corporation, Louisville, Kentucky). The lamp operates at 405nm with a peak irradiance of 1500mW/cm^2^ at a 50mm working distance [48]. This longer wavelength was chosen, as opposed to 365 or 380 nm, to minimize harm to the cells [8]. In addition, the distance between light source and sample was extended to 21cm, so that the light irradiance was predicted to be reduced to a maximum ~85.0 mW/cm^2^, based on data provided on the manufacturer’s website [48]. The cured rice cell-laden PEGTA-LAP hydrogel disks were cut into 3-4 pieces to fit through the mouth of a 25 mL flask. Liquid suspension rice cell culture control conditions were executed with and without exposure to the UV curing process.

Cured cell-laden PEGTA-LAP hydrogels consisted of 1 g FW and 1 mL bioink (or 1 mL media for liquid suspension culture controls), and the gels or liquid solutions were added with 9 mL NB+S medium in 25 mL shake flasks. Flasks were incubated at 28 °C, 140 rpm in the dark for up to 14 days. Smaller-scale experiments, which employed 0.1 g FW cells and 0.1 mL bioink (or 0.1 mL media for liquid suspension culture controls), were also performed with results and conclusions supportive of the larger-scale experiments (data not shown).

### Regulation of rrBChE Production

All three conditions (cells in hydrogel, cells in liquid suspension + UV, cells in liquid suspension - UV) started out in NB+S growth medium on day 0. On day 6, a medium exchange was performed to remove the NB+S growth medium and replace it with sugar-free NB-S expression medium to activate the Ramy3D promoter and initiate rice recombinant butyrylcholinesterase (rrBChE) production. Spent NB+S medium in the 25 mL flask was removed by pipette, following ~5 minutes gravity sedimentation for the liquid suspension culture controls. Fresh sugar-free NB-S medium was pipetted in to achieve the same working volume. A second medium exchange, from NB-S medium to NB+S medium, was performed on day 12 to halt rrBChE production and return the cultures to growth phase for two additional days until culture termination on day 14.

Cell and supernatant samples were collected on days 0, 6, 12, and 14. For liquid cultures, homogenous samples were collected, and cells and media were separated after cells were allowed to gravity settle for ~5 minutes. For flasks with hydrogel disks, the hydrogels were collected and cut into small pieces to obtain cell samples, and the remaining media was obtained free from cells or solid fragments. Experiments were set up with multiple flasks for each cell condition, so that an entire flask could be harvested at each sample point. Two biological replicates were performed, and data are shown separately for each replicate, unless otherwise indicated. Technical triplicates were performed for every biological replicate sample, and all results report average values with ± one standard deviation error bars.

### TTC Cell Viability Assay

Cell viability was assessed by the absorbance values from an assay using 0.4% (w/v) 2,3,5-triphenyl-2H-tetrazolium chloride (TTC), measured as grams TTC per mL of 50 mM sodium phosphate buffer, pH 7.5. TTC is used to differentiate between metabolically active and inactive cells. It is a colorless compound that is enzymatically reduced to a red compound, TPF (1,3,5-triphenylformazan), in living cells due to the activity of various dehydrogenases involved in cellular respiration, but it remains in its unreacted, colorless state in dead cells. Following the TTC assay protocol, approximately 0.1 g FW of cells were transferred to a 2 mL Eppendorf tube and washed with 0.8 mL of ddH2O. For cells from suspension culture, the cell mass was weighed directly whereas for cells in hydrogel pieces, it was assumed that the solid pieces contained 50% (w/v) cells (same percent from day 0 setup), so that 0.2 g of hydrogel with immobilized cells were used for the assay. After the cells were washed and water was removed, 0.8 mL of TTC was added. The tubes were immediately kept in the dark and incubated for 12-24 hours at room temperature without shaking. By the end of this time, the viable cells had all turned bright red, and pictures were taken. Then, the TTC solution was carefully removed, and the cells were washed with 0.8 mL of ddH_2_O. The water was taken off, and 1.5 mL of 95% reagent ethanol was added to extract the TPF into the supernatant. During the extraction, the tubes were kept at 60°C in a heating block, for 2-24 hours. The minimum time for the red color to be completely extracted from the cells was 2 hours, but no adverse effect is observed in continuing the extraction overnight. After extraction, the supernatant was removed (no centrifugation necessary), triplicate absorbance readings were taken per sample at 485 nm on a SpectraMax 340PC spectrophotometer (Molecular Devices, San Jose, CA) and the absorbance values were normalized by g FW of cells in the extraction tubes. The normalized absorbance values give an approximate indication of the viability of cells in the sample.

### Fluorescent Measurement and Confocal Microscopy for Cell Viability

Cell viability was also assessed, in a separate study, using confocal microscopy. The cells in hydrogel were prepared by pipetting 50 μL of 50% (w/v) rice cells in bioink, made of 12% (w/v) PEGTA and 0.1% (w/v) LAP, into PDMS mold disks, which were temporarily attached to glass slides. The samples were then cured under UV light at 405 nm, 100 mW/cm^2^ for 10 seconds, at 12.5 cm distance between light and sample. The immobilized cell samples were placed in hanging cell culture inserts with 0.4 μm pore size. The inserts were suspended in 24-well culture plates with 800 μL of media at the bottom of the wells to hydrate the gels. For the control conditions of cells in liquid media with or without UV curing, the 50 μL solid discs were replaced with 50 μL of cell suspension, also at 50% (w/v) cell loading. Just as in the larger-scale studies, cells started out in NB+S medium on day 0, were transferred to NB-S medium on day 6, and were put back into NB+S medium on day 12. On days 0, 6, 12, and 14, cell samples were collected via a 300 μL pipette with the tip cut off and were transferred to 24-well black, glass-bottom imaging plates (Eppendorf AG, Hamburg, Germany) to conduct the viability assay. Cells were stained using a standard LIVE/DEAD BacLight Bacterial Viability Kit (L7012, Thermo Fisher Scientific, Waltham, MA), and fluorescence signals were measured using a Synergy H1 Hybrid plate reader (BioTex, Winooski, VT). The green channel measures viable cell signal with SYTO 9 stain (excitation/emission: 470/540 nm) while the red channel measures non-viable cell signal with propidium iodide (excitation/emission: 470/620 nm). Viability of the cells was measured for 14 days, and the ratio of live to dead signal was recorded over time. The stained cells were imaged using a Zeiss LSM 700 confocal microscope (Zeiss, Oberkochen, Germany).

### Sugar Analysis

Sucrose and glucose levels from the media in each sample were measured with the YSI 2900 Biochemistry Analyzer (Xylem, Inc., Rye Brook, NY). Fresh NB+S medium contains 30 g/L sucrose and no glucose as prepared, though measurements taken on the media just after inoculation were closer to 21 g/L sucrose and 1 g/L glucose, likely due to some of the sugar degrading during autoclave sterilization of the media. During metabolism, the rice cells convert sucrose to glucose and fructose; the rate of sucrose consumption, glucose production, and then subsequent glucose consumption are determined by the combined effects of cell metabolism and mass transfer within the hydrogel and cell aggregates for the immobilized cells.

### rrBChE Activity in Media and Crude Cell Extract

Active recombinant BChE levels from rice cells were measured with a modified Ellman assay [49]. Ellman’s substrate is made up of 0.5 mM of S-butyrylthiocholine iodide (BTCH) and 0.267 mM of 5,5′-dithiobis-2-nitrobenzoic acid (DTNB) in 100 mM sodium phosphate buffer, pH 7.4 [38]. BChE reacts with BTCH to give thiocholine and butyrate. Thiocholine further reacts with DTNB to yield TNB. In aqueous solution at neutral and alkaline pH, TNB ionizes and generates a yellow color. The color production is quantified during the kinetic assay via absorbance measurements.

The Ellman assay was performed on both media and crude cell extract to determine the secreted and cell-associated levels of BChE, respectively, from each sample. To obtain the crude cell extract from cells grown in suspension, approximately 0.9 g FW of cells (as determined by the initial total cell g FW minus the cell g FW used for TTC assay) was first washed with 100 mM sodium phosphate buffer, pH 7.4 at an approximate 1:1 cell:buffer ratio, in 5 mL Eppendorf tubes or 15mL Falcon tubes. The samples were centrifuged down, and the supernatant was removed. Then, the samples were resuspended in 1 mL of cold (4°C) 100 mM sodium phosphate + 100 mM NaCl buffer, pH 7.4. The cells were immediately homogenized with a Tissue Tearor Model 985370 (Biospec Products, Bartlesville, Oklahoma) at maximum speed for 60-90 seconds. The cells embedded in the hydrogel were more difficult to homogenize, and it took ~180 seconds to crush both the hydrogel and the cells. The homogenized samples were centrifuged again for a minimum of 10 minutes, and the supernatant containing rrBChE was transferred to a new tube for further tests. Tubes of crude extract and of media were stored at 4°C or −20°C for short-term use or at −80°C for long-term storage.

The Ellman assay for each media sample and crude extract sample was done in triplicate in a 96-well plate. First, samples were diluted, as necessary, with 100 mM sodium phosphate buffer, pH 7.4, such that they were between 200-800 mOD/min. Then, 50 μL of sample was placed in each well, and as soon as 150 μL of Ellman’s substrate was added, the plate was put into a SpectraMax 340PC spectrophotometer (Molecular Devices, Sunnyvale, CA), which read the absorbance at 405 nm for 3-5 minutes with readings every 15 seconds, at 25°C. In this assay, human BChE (hBChE) and previously purified rrBChE were used as positive controls, and sodium phosphate buffer was used as a negative control. To calculate the amount of rrBChE in the sample, as the concentration in μg/mL or μg/g FW, from units of activity, the specific activity of crude plant-made BChE was assumed to be 260 U/mg [3,38].

### Total Soluble Protein in Media and Crude Cell Extract

The concentration of total soluble protein (TSP) from rice cells was measured with the Bradford assay [50]. The assay involves binding of protein to Coomassie Brilliant Blue G-250, which then causes the absorption peak of the dye to shift from 465 nm to 595 nm. The total protein in the sample is determined from the absorbance reading. In this analysis, bovine serum albumin (BSA) solutions at 0.5, 0.4, 0.3, 0.2, 0.1, and 0 mg/mL concentrations were used as standards. The standard curve was generated with a quadratic or linear fit (whichever had higher R^2^ value) to map absorbance values to concentration values. Rice cell extract samples were diluted 10-20 times with 100mM sodium phosphate, pH 7.4 buffer to ensure that absorbance readings were within range of the standard curve while media samples did not require any dilution. To perform the assay, 10 μL of each standard or sample was added to 3 wells of a 96-well plate, followed by 190 μL of 1x Bradford reagent. After 2 minutes, the plate was placed in a SpectraMax 340PC, spectrophotometer (Molecular Devices, Sunnyvale, CA) and the absorbance determined at 595 nm. This absorbance was converted to TSP using the equation fit to the standard curve.

### rrBChE Characterization

SDS-PAGE and Western Blot assays were performed to characterize the protein content of the media and cell extracts. Previously purified rrBChE was used as the positive control. Samples were combined with 5% (v/v) β-mercaptoethanol to reduce disulfide bonds in the tetramerized rrBChE protein into 65 kD monomer subunits. Samples were then combined with 4x Laemmli buffer and heat denatured at 95°C for 5 minutes in a heating block. Next, 35 μL of each sample was loaded onto a BioRad Mini-PROTEAN TGX Stain-Free Precast gel along with 8 μL of each BioRad Precision Plus Protein Ladder (Unstained and All Blue) in separate lanes, and the gel was run at 200 V for 35-40 minutes. The gel was imaged on a BioRad ChemiDoc system (Hercules, CA) with the stain free gel protocol and then transferred to a Biorad Trans-Blot Turbo nitrocellulose membrane using a BioRad Trans-Blot Turbo System (Hercules, CA) with the built-in Mixed Molecular Weight protocol for 7 minutes. After the transfer was complete, the blot was washed three times (TBST solution for 5 minutes at room temperature and 60 rpm nutation) and then blocked overnight at 4°C in a 1% (w/v) casein in PBS blocking solution. The next day, the blot was washed three times again and then treated with a secondary antibody binding step (1:500 mouse anti-BChE HRP in blocking solution for 1 hour at room temperature and 60 rpm nutation). Finally, the blot was washed three times again, incubated in Clarity X chemiluminescent detection solution for 5 minutes, and imaged on the ChemiDoc using the chemiluminescent, stain free blot, and Ponceau stain protocols.

## Acknowledgements

Anika Varma would like to thank Kantharakorn Macharoen, Min Du, Seongwon Jung, and Eric Knapp for their mentorship and lab assistance.

LLNL release number LLNL-JRNL-818784.

## Supporting Information

**Figure S1.**
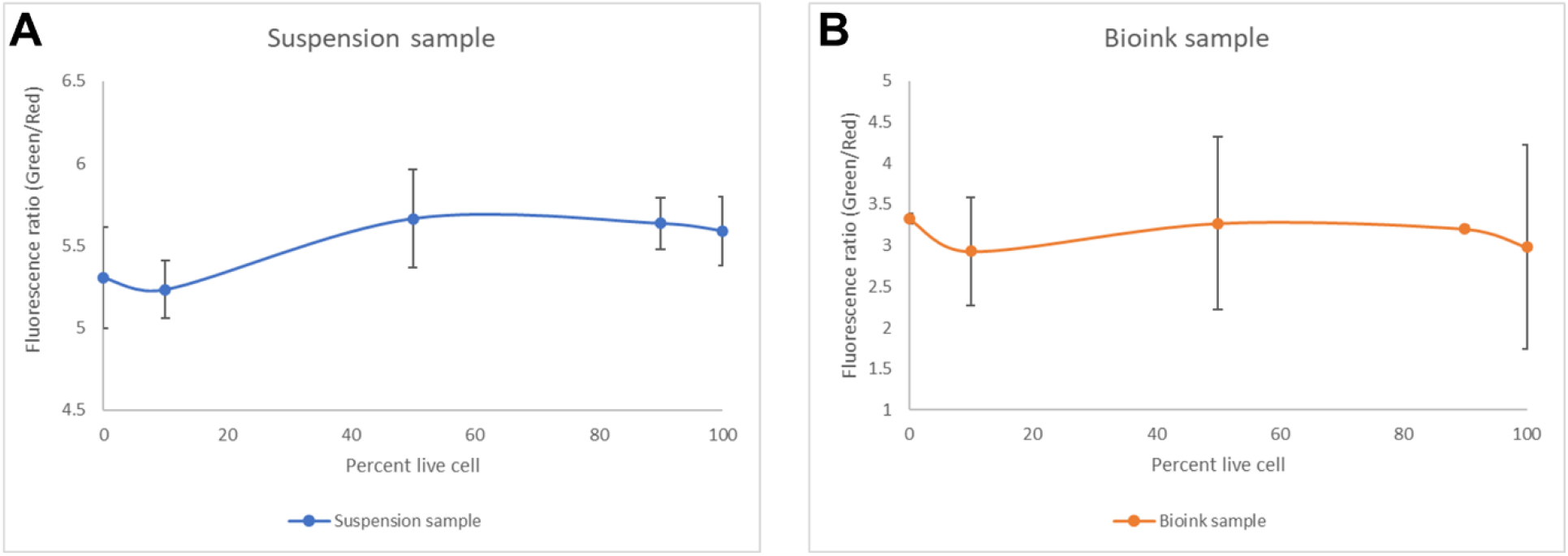
Calibration curve for fluorescence viability assay. Fluorescence (green/red) signal vs. % live transgenic rice cells stained with LIVE/DEAD BacLight Bacterial Viability Kit. (A) Calibration curve for cells in liquid suspension. (B) Calibration curve for cells immobilized in bioink (hydrogel). Error bars are ± 1 standard deviation from biological duplicates

**Figure S2.**
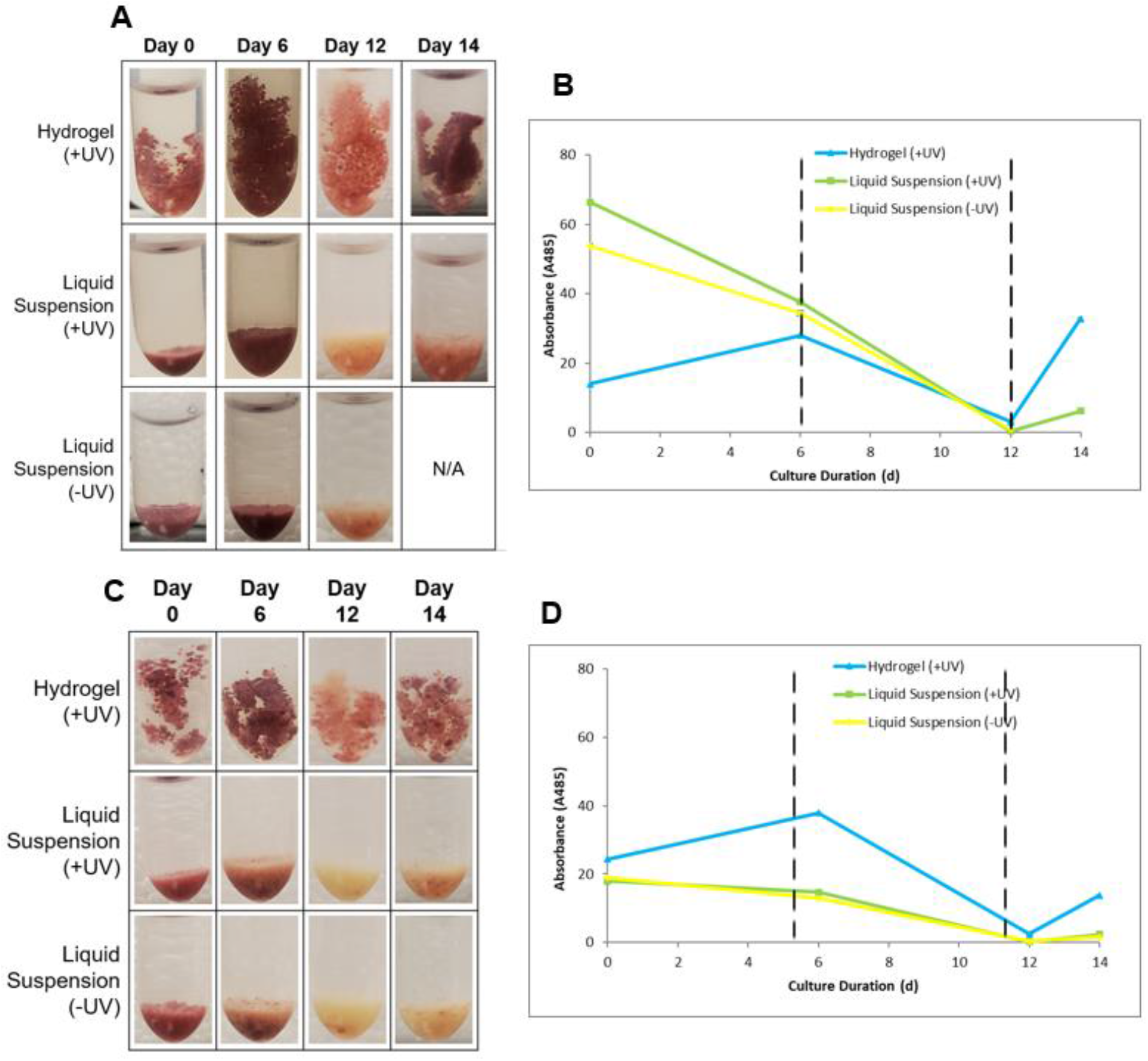
Transgenic rice cell viability from TTC assay. Cell viability over the course of 14-day culture (with media exchanges from sugar-rich to sugar-free medium on day 6 and from sugar-free to sugar-rich medium on day 12), as shown by TTC assay data separated by experiment. (A) Pictures of TTC-stained cells and (B) TTC assay absorbance readings for cells from experiment #1 (15-day old cells). (C) Pictures of TTC-stained cells and (D) TTC assay absorbance readings for cells from experiment #2 (33-day old and 13-day old cells). Error bars on absorbance graphs represent ± 1 standard deviation from technical triplicates.

